# Exp2Ipynb: A general machine-learning workflow for the analysis of promoter libraries

**DOI:** 10.1101/2020.12.14.422740

**Authors:** Ulf W. Liebal, Sebastian Köbbing, Lars M. Blank

## Abstract

Strain engineering in biotechnology modifies metabolic pathways in microorganisms to overproduce target metabolites. To modify metabolic pathway activity in bacteria, gene expression is an effective and easy manipulated process, specifically the promoter sequence recognized by sigma factors. Promoter libraries are generated to scan the expression activity of different promoter sequences and to identify sequence positions that predict activity. To maximize information retrieval, a well-designed experimental setup is required. We present a computational workflow to analyse promoter libraries; by applying this workflow to seven libraries, we aim to identify critical design principles. The workflow is based on a Python Jupyter Notebook and covers the following steps: (i) statistical sequence analysis, (ii) sequence-input to expression-output predictions, (iii) estimator performance evaluation, and (iv) new sequence prediction with defined activity. The workflow can process multiple promoter libraries, across species or reporter proteins, and classify or regress expression activity. The strongest predictions in the sample libraries were achieved when the promoters in the library were recognized by a single sigma factor and a unique reporter system. A tradeoff between sample size and sequence diversity reduces prediction quality, and we present a relationship to estimate the minimum sample size. The workflow guides the user through analysis and machine-learning training, is open source and easily adaptable to include alternative machine-learning strategies and to process sequence libraries from other expression-related problems. The workflow is a contribution to increase insight to the growing application of high-throughput experiments and provides support for efficient strain engineering.

**Availability:** Freely available on GitHub: https://qithub.com/iAMB-RWTH-Aachen/Exp2Ipynb and licensed under the terms of GPLv3.

**Supplementary information:** Supplementary data available in the Git folder.

## 1 Introduction

Metabolic engineering aims at optimizing metabolite production by adjusting the activity of native and heterologous enzymes. A frequently manipulated factor of activity is the enzyme concentration which can be regulated on the transcriptional, translational and post-translational level. For each level, computational models were developed allowing realistic simulations and supporting rational adjustment. Models of σ^E^ transcription in *E. coli* were developed by Rhodius *et al.* (2010; 2012) and allow analysis of the effects of upstream regulating elements and core promoter. Translational components affect enzyme concentration via ribosome binding site, codon usage and mRNA stability. The effect of the ribosome binding site depends on the homology of the Shine-Dalgarno sequence and mRNA secondary structures and for *E. coli* a predictive model was published (Salis *et al.*, 2009) and recently integrated with a resource allocation model (Nóbel and Picó, 2020). The codon usage, mRNA stability and enzyme concentration are correlated and jointly determine enzyme concentration in *E. coli* (Kudla *et al.*, 2009; Boel *et al.*, 2016), and a predictive model of mRNA stability was published by Cetnar & Salis (2020). Finally, protein stability determines concentration and a review of predictive models is given by Fang (2020). Although the models have reported high correlation between prediction and experiment, the massive evaluation of 244.000 sequences for translational effects during σ^70^ expression in *E. coli* achieved lower correlation (Cambray *et al.*, 2018). In conclusion, the relative impact of expression factors is hard to estimate on the genome level.

The promoter sequence is important for strain engineering because it allows fine-tuning and modular exchange of elements. The composition and regulation of promoters have been intensively studied (Kalisky *et al.*, 2007; Zaslaver *et al.*, 2009; Kochanowski *et al.*, 2017). For example, Rhodius et al. (2010, 2012) investigated the expression strength of 60 σ^E^ promoters and analyzed the impact of promoter boxes and upstream regulating elements on expression. A modular promoter system was developed by Mutalik *et al.* (2013), that reduces interference from the sequences of the gene of interest, resulting in reproducible promoter activities. Promoter libraries are routinely constructed to generate a set of promoters with defined activities generated to provide reference expression activities (Jensen *et al.*, 1993; Alper *et al.*, 2005; Hammer *et al.*, 2006; Balzer *et al.*, 2013; Köbbing *et al.*, 2020). For example, Meng *et al.* (2013, 2017) analyzed 98 σ^70^ promoter sequences and fine-tuned heterologous expression with predicted synthetic promoters. The same system was analyzed by Zhao et al. (2020) with over 3500 promoter sequences. In *Bacillus subtilis* Liu *et al.* (2018) employed a synthetic promoter library with 214 sequences to fine-tune pathway activity for metabolite overproduction. To facilitate metabolic engineering in *Geobacillus thermoglucosidasius,* a library with 80 sequences was tested for both the expression of GFP and mOrange (Gilman *et al.*, 2019). These libraries have been generated with a sample size to implement model-based sequence analysis and rational promoter development.

Cross-evaluation of promoter library analysis is impeded because of different tools and reporting standards. To unify the analysis and to provide a platform suitable for easy reconfiguration, we have developed a general workflow for promoter library analysis called the *Exp2Ipynb.* The *Exp2Ipynb* is a Python Jupyter Notebooks and consists of workflows to (i) generate a statistical overview to sequence and activity, (ii) train machine-learning algorithms for prediction and extraction of feature importance, (iii) evaluate the performance of the estimator, and (iv) to predict new sequences with defined activity. We tested the workflow on an existing promoter library for σ^70^ in *Pseudomonas putida* KT2440, expanded by new measurements. The results revealed limitations of one-factor-at-a-time experimental designs for machine learning. Subsequently, we used six published libraries to compare their composition and the resulting machine learning performance.

## 2 Workflow and implementation / Methods

We constructed a workflow for the statistical analysis and training of machine learning models of genomic sequence data called *Exp2Ipynb.* The workflow is available as a collection of Jupyter Python Notebooks and follows the guidelines for code development (Rule *et al.*, 2019). The workflow is freely available via Git and can be easily modified. The Notebooks represent the following steps: (i) analysis of statistical properties of sequence and reporter, (ii) training of the machine learning algorithm, (iii) evaluation of prediction performance, (iv) prediction of new sequence with defined activity. The input to the machine-learning estimator are sequences and the output is a list of associated activities.

### 2.1 Data preparation and statistical analysis

A configuration file stores globally used variables that are adjusted to each analysis. The data input is a comma-separated-value file (csv) with at least an identifier column, a sequence column and a response column, with header names defined in the configuration file. The sequence column only accepts DNA abbreviations (A, C, G, T) with an identical length. The output file names and figure file types can be defined. The notebook ‘0-Workflow’ guides through the construction of the configuration file.

The statistical analysis is conducted in the notebook ‘1-Statistical-Analysis’. The libraries may contain activity measurements of replicates or mean and standard deviation. Outliers in the original data set can be removed from further analysis. Machine learning performance is improved if replicates are explicitly available and the workflow enables the re-generation of replicates based on mean and standard deviation. The approach is valid for normal-distributed data and adds a reasonable prediction bias. The feature list is generated from a one-hot encoding of the sequence, plus the overall GC content.

The statistical analysis provides an overview to the following metrics:

- sequence diversity
- position diversity
- nucleotide-position expression statistics
- expression strength distribution
- cross library expression strength

The promoter diversity is evaluated by the sequence and position diversity. The sequence diversity represents the coverage of the sequence sample space and is calculated as the amount of nucleotide differences over the sequence relative to a reference sequence or between all sequences, and ranges between ‘0’, identical to ‘1’ all nucleotides are different. The reference sequence can be provided with the configuration file, or it is generated automatically by finding the most common nucleotide on each position. For large libraries >10^3^ samples, the full pairwise distance is costly to compute. The position diversity informs about how many nucleotides have been sampled for each position. It is visualized with two bar plots: (i) cumulative number of each nucleotide tested on each position, and (ii) the position specific entropy (*H*’), reflecting the information content at each position (*i*), along the sequence (*R*) with the position-related proportional abundance of nucleotides (*p_i_*):

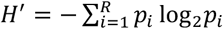

The expression statistics for all nucleotides and positions informs how each nucleotide contributes to the expression. Each sequence is transformed into a matrix with rows as position and four columns, representing the one-hot encoding of the nucleotides (A, C, G, T). This sequence-one-hot matrix is multiplied by the expression strength, resulting in a matrix with the expression strength at correct nucleotide-positions in the matrix. The matrices yield the expression average and variance on each nucleotide position and indicates which positions are associated with higher or lower expression. The expression strength is visualized with a histogram and a scatter plot for cross library expression.

### 2.2 Predictor training and performance

The workflow supports classification and regression. The advantage of regression is the prediction of quantitative expression values for sequences, but the training requires high data quality (sufficient sample size and position entropy). Classification provides qualitative predictions *(e.g.,* low-medium-high), with stronger predictions even for small data sets. The implemented machine learning routines are random forest (RF), gradient boosting (GB), and support vectors (SV). The input features are nucleotides on each position and the overall sequence GC-content, while the predicted target variable is the expression strength. Because the input features can be high-dimensional depending on the sequence length, additional feature selection procedures are implemented.

The performance of the predictions can be improved by feature selection and shifting from regression to classification. The entropy in each position of the input sequence reflects the diversity of tested nucleotides and typically, sound predictions can only be made if the entropy is high enough. Note, however, that non-canonical nucleotides in conserved promoter (box)-regions will result in drastic expression deficiency. Thus, a low diversity can also reflect strong predictive power where sequence diversity is experimentally inaccessible. A cut-off can be chosen in the configuration file to exclude positions below a defined entropy.

The choice of classification or regression is controlled by the parameter ‘Response_Value’ in the configuration file. If the ‘Response_Value’ is >1, the output is binned in the number of bins according to the value. If the ‘Response_Value’=1, the original activity values are used as prediction targets and if the ‘Response_Value’=0, the expression data is centered with zero mean and unit variance. The bins are generated as equal sized buckets (python pandas ‘qcut’) and the bin label is used as the target prediction. Data centering is a requirement for using support vector machines. The data is split into training and test set, with a default ratio of 9:1 and a grid search on the training data identifies the optimal hyperparameter.

The performance evaluation provides metrics for correlating the experimental and predicted outcomes and informs about important features for tree-based methods. The performance evaluation is based on cross-validation with 9:1 data separation that moves identical sequences (i.e. replicates) consistently using group shuffle split to the test or the training set. The group shuffle split ensures that the test sequences are unknown to the regressor. The performance evaluation is based on the R2-(regression) and weighted F1-score (classification) for training set and the test set. The prediction uncertainty metric is determined by the coefficient of variance of the mean R2 and F1 prediction over the training set cross validation. The tree-based methods allow the extraction of feature importance and represent the contributions of each nucleotide-position to the prediction. They are extracted and visualized with a Logo-plot (Tareen and Kinney, 2019).

### 2.3 Synthetic sequence characterization

The workflow allows to identify the exploration space covered by the samples for which reliable predictions are possible and to test novel sequences within the exploration space. The exploration space spans the sequences for which reasonable predictions can be expected and is bounded by the sequence and position diversity. Only for sufficiently sampled nucleotides at any position can the regressor derive activity rules and predict reasonable activities. In addition, the sequence distance of an unknown sequence with respect to the reference sequence of all other sequences should be lower than the sequence distances in the training samples because unknown regulatory or physiological events can interfere with sequences deviating too much from the samples. The user defines the cut-off for the exploration space based on the visualizations of position and sequence diversities in the statistical analysis. The sequences of the exploration space can be generated exhaustively or a random subset of defined size, and results in a synthetic promoter library with predicted expression activities. The workflow allows to select synthetic sequences close to a target expression and the synthetic library can be statistically analysed similarly to the experimental library.

### 2.4 Experimental library construction

Construction of the single nucleotide polymorphism (SNP) library was done by PCR with plasmid pBG14g as template and different overhanging oligonucleotides containing single degenerate nucleotides inside the P14g promoter sequence (Zobel *et al.*, 2015). Generated spacer fragments and vector pBG were digested with *Pac*I and *Avr*II (New England Biolabs) at 37°C. Digested backbone and promoter containing PCR fragments were ligated with T4 ligase (New England Biolabs) at room temperature for 30 minutes. Transformation into chemically competent *Escherichia coli* PIR2 cells was done by heat shock (Hanahan, 1983). Further characterization of plasmid bearing *E. coli* PIR2 cells was based on msfGFP fluorescence measurements with a synergyMX plate reader (Biotek) at an excitation wavelength of 488 nm and emission wavelength of 520 nm. Plasmids containing different synthetic promoter sequences were sequenced (Eurofins Genomics) and afterwards genomically integrated into the *attTn7* site of *P. putida* KT2440 by mating (Zobel *et al.*, 2015). For promoter characterization in *P. putida* KT2440 a Biolector was used (M2P Labs). Cells were grown in minimal medium as described by Hartmans *et al.* (1989) with 20 mM glucose as carbon source. The Biolector system measured optical density at 620 nm, msfGFP was measured at excitation wavelength at 488 nm and emission wavelength of 520 nm. Scattered light was correlated to OD_600_ with a dilution series of a stationary phase culture. Promoter activity is reflecting the slope of the function of fluorescence over OD_600_ at the beginning of the exponential phase. More detailed information is given in Köbbing et al. (2020).

## 3 Results

The workflow was first applied on a newly extended *Pseudomonas putida* KT2440 synthetic promoter library, followed by a cross analysis of six published libraries. The analysis of our *P. putida* KT2440 library will show how the workflow can be used, and we will highlight shortcomings of the experimental design for machinelearning driven analysis. In the cross-library comparison, we investigated how the different library parameters of sample size, diversity and feature number impact the predictions.

### 3.1 *P. putida* single library analysis

We used the workflow to analyse a synthetic library in *P. putida* KT2440 driven by σ^70^-dependent promoters and measured with GFP published previously (Köbbing *et al.*, 2020) but expanded by eight new sequences. The goal was to identify sequence features responsible for expression strength. Experimentally, the sequence diversity was generated by single nucleotide exchanges in 28 nucleotides upstream of the transcription start site resulting in 63 unique sequences. For the analysis we used 40 nucleotides of the promoter and used three approximately equal sized bins for the activity classification. The sequences had low information content on most positions (Figure 1 (A) and (B)) allowing only predictions of categorized expression values, a regression predicted not better than random (not shown).

**Figure 1:**
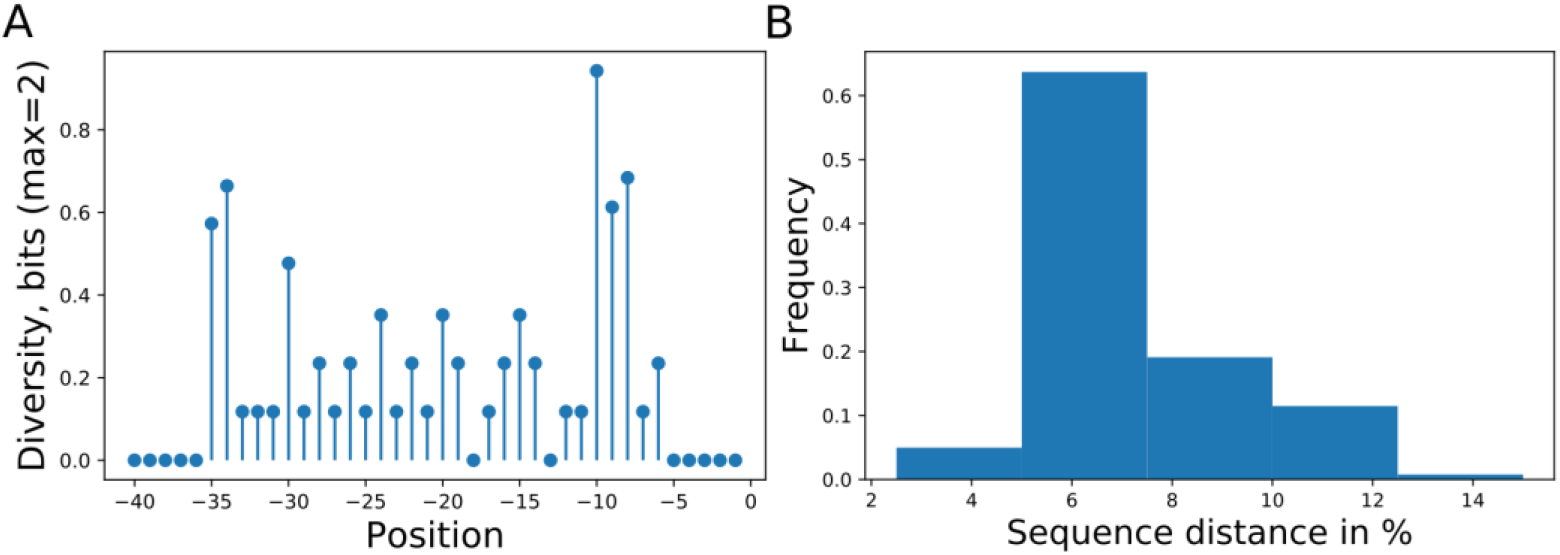
Promoter sampling diversity of the complete data set for nucleotide variations on each sequence position (A) and mutual sequence distances (B). The promoter library is based on Köbbing et al. (2020) with additional sequences totaling 63 different promoters with mutations directly upstream of the transcription start site.

A classification estimator can predict the approximate magnitude of expression. A reasonable estimation can only be performed on features with sufficient information content, and positions with a higher entropy than 0.2 bits were included for prediction. Figure 1 (A) shows the entropy of the full data set, and because the cut-off is applied to the training subset, the following positions are additionally neglected [-6,-15-20,-22,-26]. Decreasing the entropy threshold had no effect on the performance (not shown). The classification details are given in Table 1 and the high F1-score variation indicates that the low nucleotide redundancy on each position affected sample separation during cross validation. The detailed prediction results of the training and test set are shown as confusion matrix in Figure 2 (A). The GC-content is among the most important features for the prediction probably because it has the highest entropy of all features (1.2 bits). Along with the GC-content, the strongest determinant of expression was the nucleotide ‘T’ at position –35 and −34, corresponding to the fact that a nucleotide exchange at these positions cause a strong expression decrease (Figure 2 (B)).

**Figure 2:**
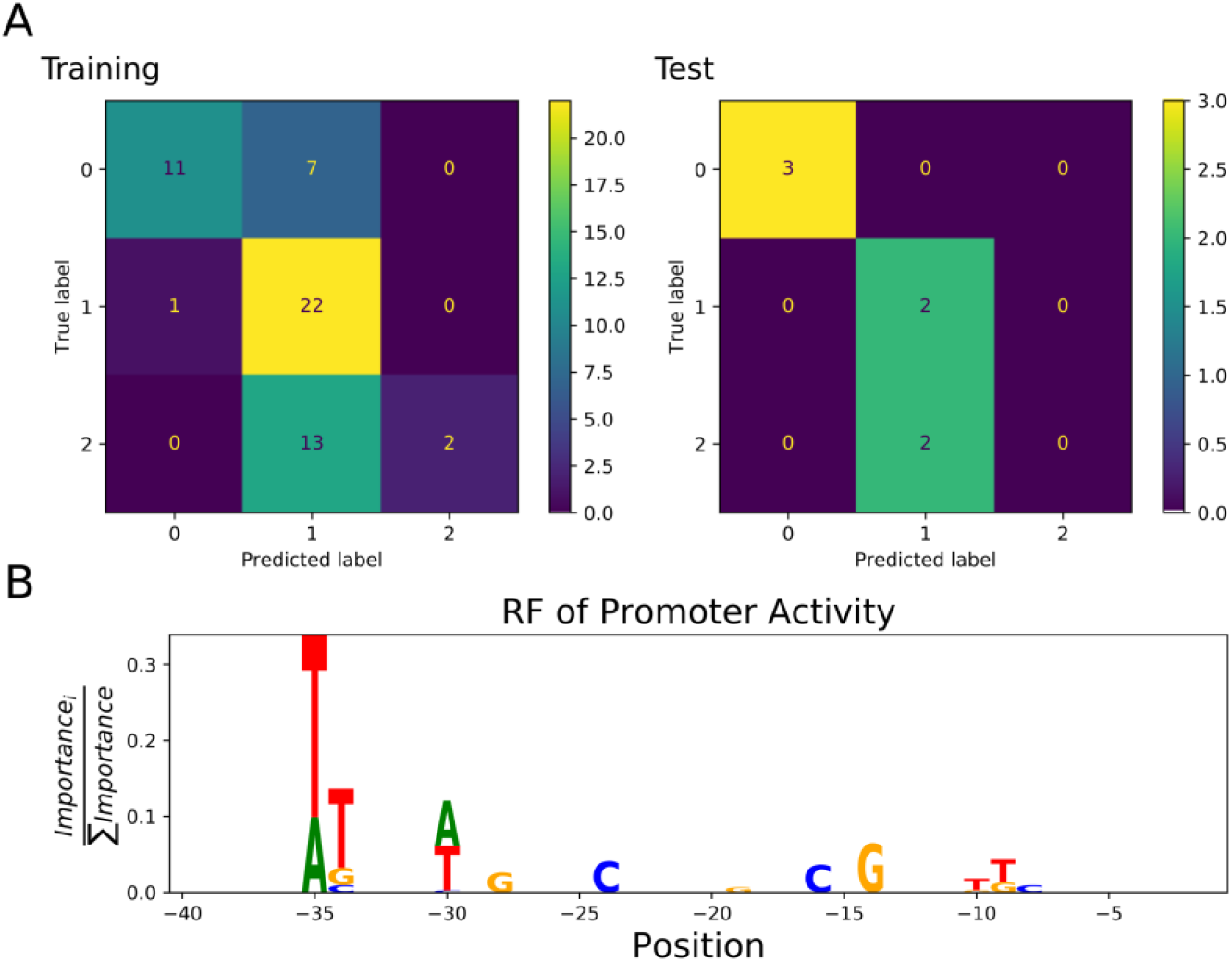
Confusion matrix showing classification quality for Training (n=49) and Test (n=6) sets (A). The classification labels are ‘0’ for low (0.4-18.7 GFP), ‘1 ‘ for medium (18.7-25 GFP), and ‘2 ‘ for strong expression (25-35 GFP) (Data source Köbbing et al., 2020). Sequence logo of positions used by the RF to predict expression activity class (B). The –35 box was dominating the prediction whereas the –10 box had lesser importance on prediction despite similar high diversity (Figure 1).

**Table 1:** Classification quality report for random forest (RFC), gradient boosting (GBC) and support vector (SVC). The training was performed with the same Train-Test (10% hold out) and the entropy threshold was 0.2 bits resulting in five positions as input features. The weighted f1-score represents prediction accuracy and is calculated from precision and recall. The feature importance for nucleotide-position and GC-content was only determined for tree-methods (RFC and GBC).

|  | RFC | GBC | SVC |
| --- | --- | --- | --- |
| Run time (s) | 140 | 599 | 25 |
| CV: F1-score | 0.44 (+/-0.41) | 0.50(+/-0.41) | 0.47(+/-0.42) |
| Train: F1-score | 0.58 | 0.89 | 0.88 |
| Test: F1-score | 0.62 | 0.46 | 0.14 |
| GC-Content | 2<sup>nd</sup> | 1<sup>st</sup> | N.A. |
| Top 3 FI | -35:T:0.24<br>-34:T:0.1<br>-35:A:0.1 | -34:T:0.08<br>-35:A:0.05<br>-14:G:0.05 | N.A. |

### 3.2 Cross-library analysis

In the following, six published bacterial promoter libraries were analyzed with the aim to highlight the effect of the library setup on the ML estimation quality. The data is available online in the *Exp2Ipynb* package at GitHub. The libraries were measured in *E. coli*, *Geobacter thermoglucosidasius*, and *B. subtilis* and were targeted to specific or unknown sigma factors. All libraries differ from each other in terms of combinations of library sample size, tested sequence length and sequence diversity (Table 2). We evaluated how each parameter affects estimation quality.

**Table 2:** Details of the six published libraries used to compare estimation quality in response to library experimental design. The Avg.Seq.Dist’ represents the average distance of all sequences to a reference sequence that is composed of the most common nucleotide at each position..

| Reference | ID | Organism | Transcr. Factor | Reporter | Run-Time (s) | # Samples | # Features | Avg.Seq.Dist. |
| --- | --- | --- | --- | --- | --- | --- | --- | --- |
| This Work | a | *P. putida* | $\sigma^{70}$ | GFP | 140 | 63 | 61 | 0.06 |
| Rhodius et al. (2010) | b | *E. coli* | $\sigma^E$ | various | 134 | 59 | 121 | 0.61 |
| Meng et al. (2013) | c | *E. coli* | $\sigma^{70}$ | GFP | 143 | 98 | 409 | 0.16 |
| Zhao et al. (2020) | d | *E. coli* | $\sigma^{70}$ | GFP | 476 | 3543 | 289 | 0.09 |
| Gilman et al. (2019) | e | *G. therm.* | N.A. | GFP | 138 | 81 | 397 | 0.69 |
| Gilman et al. (2019) | f | *G. therm.* | N.A. | mOrange | 145 | 81 | 397 | 0.69 |
| Liu et al. (2018) | g | *B. subtilis* | $\sigma^A$ | GFP | 144 | 206 | 105 | 0.35 |

Three values are important for analysis of promoter libraries: (i) the fidelity of gene expression prediction, computed by the F1-score (see methods), (ii) the confidence of the prediction, computed by the coefficient of variation, and (iii) important sequence features for expression, computed by the feature importance of random forest and gradient boosting strategies. The analysis was based on a classification task with three classes: low (‘1’), medium (‘2’), and high expression (‘3’). The analysis was performed on the original data of the promoter libraries without detailed processing. For quality assessment, we generated three additionally data sets based on partitioned sequence regions from the Meng *et al.* (2013) and the Zhao *et al.* (2020) data sets. The original sequence from Meng *et al.* (2013) is 224 nucleotides (nt) long and we extracted the starting 40 nt (‘h’) and last 40 nt (‘i’), assuming that few predictive positions are contained in the new sequences. Zhao *et al.* (2020) tested 113 nt ranging from upstream regulating elements down to the coding sequence. Again the first 40 nt (‘k’) were extracted, corresponding to the upstream regulating element.

The predictions were generally weak even with large data sets. The best predictive quality was achieved by the promoter library with more than 3500 sequences (Figure 3, (A), library ‘d’). However, the high prediction quality is based on a low sequence diversity with 0.1 (see methods). The F1-score is independent of the number of tested input features (Figure 3 (A)). It is important to judge the stability of the predictions via cross validation as determined with the coefficient of variation of the weighted F1-score (F1-CoV). The training sample size has a hyperbolic decreasing effect on the F1-CoV (Figure 3 (C)): below 500 samples the F1-score depends on sample composition and is predicted unreliably (‘a’, ‘b’, ‘c’ and ‘e’, ‘f’, ‘g’). As expected, the higher the sequence diversity, the more variable was the F1-score, again strongly depending on sample composition (‘b’ and ‘e’, ‘f’) (Figure 3 (D)). Libraries ‘a’ and ‘c’ display a notable position with a high F1-CoV despite low sequence diversity, caused by low promoter library size. An important information is the feature importance that informs about critical drivers of expression. Figure 3 (B) indicates how the number of features affects the sum of the three top important features derived from the RF. The top-three feature importance sum decreases linearly with increasing numbers of features.

**Figure 3:**
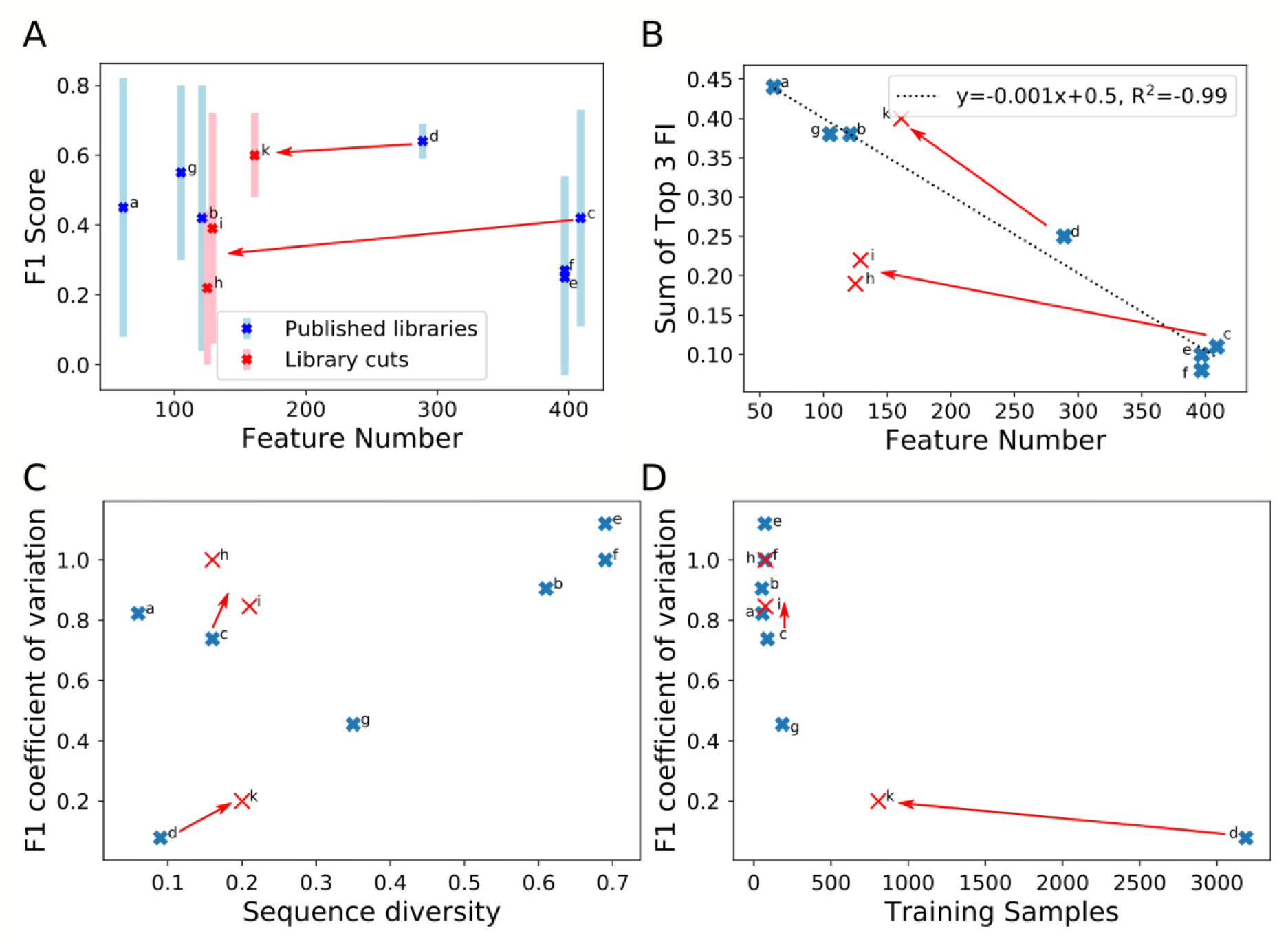
Comparison of library properties on classification quality parameters for an estimation using RF. The estimations were performed on the training set of each library with 9:1 cross validation. The seven full length promoter libraries are listed in Table 2. Moreover, sub-sequences of 40nt were extracted from the promoter start (‘h ‘) and end (‘i’) of Meng et al. (2013) (‘c’) and start (‘k’) of Zhao et al. (2020) (‘d’). F1-score (A) and the sum of the top three features (B) in response to the feature number. The coefficient of variation of the F1-scores in response to sequence diversity (C), and number of training samples (D).

We were interested to examine how the performance is affected when subsequences were extracted from libraries. We chose the libraries from Meng *et al.* (2013) and Zhao *et al.* (2020) because they report high feature number with low sequence diversity. The starting (‘h’) and ending (‘i’) 40 nt were extracted from the promoter sequence in Meng *et al.* (2013), which had low features with importance. The rational to use the starting 40 nt from Zhao *et al.* (2020) (‘k’) is that they contained the upstream promoting element. The sequence extraction reduces the sample size because more redundant sequences are generated, and sequence diversity increased because nucleotide diversity covers a shorter sequence. Figure 3 shows that the partitioned dataset ‘k’ (from Zhao *et al.* (2020)) maintained the quality of prediction from the original dataset: the F1-CoV and top three feature contribution increased proportionally to the original data (B, C and D). This proportional movement was not displayed by the extracted sequences (‘h’, ‘i’) from Meng *et al.* (2013). Thus, sequence position impacting expression were identified in the upstream regulating elements of Zhao *et al.* (2020), but not in the first and last 40 nt of the sequences of Meng *et al.* (2013). The examined libraries form a triangle in prediction reproducibility versus sequence diversity graph (Figure 3 (C)): increase of sequence diversity directly reduced prediction reproducibility, unless the sample size was critically reduced (data sets ‘a’+’c’). We estimated the critical data set size based on the synthetic data set ‘k’: with 800 samples on a diversity of 0.2, the sample size was still saturated, and the prediction reproducibility depended directly on the sequence diversity.

## 4 Discussion and conclusion

Synthetic promoter libraries are increasingly constructed to facilitate promoter selection with defined activities. The *Exp2Ipynb* workflow supports the analysis of promoter libraries to identify the sequence information that determines expression strength. The identification is based on statistical analysis of the average nucleotide-position associated expression, as well as the training of different machine learning algorithms. Moreover, the analysis of more complex libraries with multiple readouts, like two reporter proteins, transcript and protein level, or cross host expression can be analyzed. The tool facilitates data exploration by providing a workflow from data preparation, regressor training and performance evaluation, and testing of novel sequences within the DNA sequence exploration space. The implementation in a Jupyter notebook facilitates rapid implementation and the low-level scripting allows for direct adaptation of the workflow to specific needs.

We analyzed an expression library in *P. putida* KT2440 previously published (Köbbing et al., 2020) and amended with additional data points. The diversity was generated based on a one-factor-at-a-time approach to economically identify nucleotide-sequence effects on expression (Czitrom, 1999). The strong impact of the positions −35 and −34 and the lower impact of the last half of the −10 box (TA**TAA**T, −10, −9, −8) confirmed results in the original article (Köbbing et al., 2020, Figure 4). We could not reproduce the strong effect of the first part of the −10 box (T**A**TAAT, −11) because the position was sampled below the entropy cutoff and was neglected in the analysis. The ML analysis of low diversity and low sample size data sets did not add more information than a thorough statistical analysis. To identify the magnitude of diversity and sample size libraries necessary, we conducted a cross library analysis.

An important question for the ML-based analysis of expression libraries is: how much samples are necessary to achieve a targeted prediction uncertainty? We identified two factors that affected prediction uncertainty: sample size and sequence diversity. A higher sequence diversity requires more samples to reduce prediction uncertainty. Two libraries in the collection were limited by sample size (‘a’+’c’) whereas five libraries were limited by sequence diversity (‘b’,’d’,’e’,’f’). We found that 100 samples still resulted in a high uncertainty for sequences with an average of 20% nucleotide difference (‘a’, ‘c’), whereas 800 samples proved sufficient (‘k’). With higher nucleotide differences of 35%, 200 samples were insufficient to cover the diversity (‘g’). Other factors confounding the sequence diversity-sample size relation are effects from the heterogeneous reporter system (‘b’), or unknown sigma factor specificity (‘e’, ‘f’). More libraries are necessary to narrow the required sample size over the whole sequence diversity spectrum. Particularly libraries were missing with high sequence diversity and high sample size (Cambray *et al.*, 2018; Cuperus *et al.*, 2017).

The test sets had a low weighted F1-score in the cross-validation prediction of the classification ML-estimator. The machine learning analysis approaches of the published libraries used optimized train-test set partitions for evaluation and do not report prediction uncertainty based on cross validation. Gilman et al. calculate at a stronger correlation for their model, but when the authors tested new data, the prediction qualities were similar to those we observed. Also, the convolutional neural net trained with 500,000 sequences by Cuperus et al. (2017) achieves correlation coefficients of 0.62. The regulation of gene expression is controlled by multiple factors. Additional feature engineering can be used to increase predictability: GC-moving window, secondary structure for UTR. However, because gene expression relies on various mechanisms, there will be no single machine learning tool to fit all: machine-learning ensembles are needed, composed of different approaches tailored to the various mechanisms.

So far, the analysis of promoter libraries was conducted with scripts tailored to the data, obfuscating reproducibility and interpretability. The *Exp2Ipynb* is a contribution to harmonize analytical workflows and to enable an easy start for new groups with promoter libraries. Other toolboxes exist to support machine learning like *tpot,* GAMA, or H2O (Truong *et al.*, 2019) and more oriented towards biological data analysis like *JADBIO* (Tsamardinos *et al.*, 2020). The advantage of the *Exp2Ipynb* is to be general enough to be used across different data sets in expression studies, but remaining domain specific to facilitate simple data integration.

## Additional Material

The workflow can be downloaded from the GitHub repository: https://github.com/iAMB-RWTH-Aachen/Exp2Ipynb The repository contains all data sets used in this work.

## Acknowledgements

The authors thank Salome Nies and Dario Neves for promoter library data during initial tests. The authors are grateful for discussions with Birgitta Ebert, Tobias Alter and Bastian Kister.

